# A transcriptional activator effector of *Ustilago maydis* regulates hyperplasia in maize during pathogen-induced tumor formation

**DOI:** 10.1101/2023.03.06.531288

**Authors:** Weiliang Zuo, Jasper R. L. Depotter, Sara Christina Stolze, Hirofumi Nakagami, Gunther Doehlemann

## Abstract

*Ustilago maydis* causes common smut in maize, which is characterized by tumor formation in aerial parts of the host. Tumors result from the *de novo* cell division of highly developed bundle sheath and subsequent cell enlargement. However, the molecular mechanisms underlying tumorigenesis are still largely unknown. Here, we characterize the *U. maydis* effector Sts2 (<u>S</u>mall <u>t</u>umor on <u>s</u>eedlings 2), which promotes the division of hyperplasia tumor cells. Upon infection, Sts2 is translocated into the maize cell nucleus, where it acts as a transcriptional activator, and the transactivation activity is crucial for its virulence function. Sts2 interacts with ZmNECAP1, a yet undescribed plant transcriptional activator, and it activates the expression of several leaf developmental regulators to potentiate tumor formation. Contrary, fusion of a suppressive SRDX-motif to Sts2 causes dominant negative inhibition of tumor formation, underpinning the central role of Sts2 for tumorigenesis. Our results not only disclose the virulence mechanism of a tumorigenic effector, but also reveal the essential role of leaf developmental regulators in pathogen-induced tumor formation.

## Introduction

Plant pathogens secrete effectors to cross-talk with hosts for their benefits. To reshape the host transcriptome, some pathogens exploit effectors to directly manipulate host gene regulation in two main mechanisms. One is characterized by transcription activator-like (TAL) effectors, first described in the plant pathogenic *Xanthomonas sp*^1–3^. These effectors contain a nucleus localization signal, tandem repeat DNA binding domain and transcriptional activation domain, which can function independently as transcription factors to activate host gene expression. On the other hand, several effectors are not transcription factor themselves, but control host gene expression through interacting with host transcription factors^4^, recruit the suppressors of transcription factors^5–8^, or disrupt the assembly of transcription units^9,10^, eventually leading to the inhibition or activation of host gene expression, respectively.

*Ustilago maydis* is a fungal pathogen which causes common smut in maize. It infects all the aerial maize organs, grows locally and produces massive amounts of teliospores^11,12^. To achieve this, *U. maydis* manipulates plant cell proliferation and thereby creates additional space in the form of tumors to reside in. On maize seedling leaves, tumors consist of hyperplasia tumor cells from the *de novo* division of highly differentiated bundle sheath tissue, some of such hyperplasia cells together with mesophyll enlarged to become hypertrophy tumor cells^13^. The host mechanisms of tumor formation and the causative effectors are still unknown. Until now, the functionally characterized effectors in *U. maydis* are mainly involved in immunity inhibition^14–17^ or host metabolism manipulation^18,19^. See1 is the only effector identified that is directly involved in tumorigenesis by re-activating DNA synthesis^20^; deletion of See1 causes the inhibition of *U. maydis* induced hyperplasia cell division^13^. Different tumor cells have a very different physiology and transcriptome in between, and compared to the sourced host cells^13,21^. One therefore can hypothesize that *U. maydis* possesses effectors to directly, or indirectly manipulate the host transcriptome. In line with this, several effectors of *U. maydis* have been identified to target Topless transcription co-repressors to modulate hormonal signaling and expression of immune genes during infection^6–8^. However, little is known about how effectors modulate the host transcriptome to induce tumors. A detailed *U. maydis* transcriptome analysis revealed the temporal regulation of effector genes throughout all steps of infection^22^, and laser-captured microdissection of different tumor cells coupled with RNA-seq showed spatial, cell type specific regulation of effectors^13^. The cross-species analysis between *U. maydis* and *Sporisorium reilianum*, the closest smut relative which also infects maize but does not cause tumors, disclosed the differential regulation of effector orthologs that contribute to the distinct pathogenic development in the two species. A CRISPR-Cas9 mediated effector ortholog knock-in experiment discovered a functional diversification of an effector orthogroup UMAG_05318 - sr10075/ sr10079 during speciation^23^.

In this study, we functionally characterized the effector Sts2 (UMAG_05318, <u>S</u>mall <u>t</u>umor on <u>s</u>eedlings 2). A *U. maydis* knockout strain for Sts2 (CR-Sts2) initiates tumor formation, but the tumors fail to expand due to reduced cell division of the bundle sheath. We discovered that Sts2 is a transcriptional activator, being translocated into host cell nucleus. It interacts with ZmNECAP1, a yet unknown plant transcriptional activator, to activate the expression of leaf developmental regulators. Sts2 transcriptional activation is crucial to maintain the meristem like cell divisions which lead to hyperplasic tumor cells. Our findings disclose a tumorigenic mechanism where a *U. maydis* small effector functions as a transcriptional activator in the host nucleus to rewrite the host developmental process by hijacking the key leaf developmental regulators.

## Results

### Sts2 regulates hyperplasia tumor cell induction by *U. maydis*

We generated an open reading frame shift knockout of *UMAG_05318* using CRISPR-Cas9 mutagenesis. The resulting mutant showed similarly reduced virulence **(Supplementary Fig. 1a)** as it was described previously for a deletion strain^24^ **(Fig. 1a)**. Distinct from the typical bulged tumors caused by *U. maydis* SG200, tumors induced by the UMAG_05318 knockout mutant failed to expand in size **(Fig. 1a)**. Therefore, we named UMAG_05318 as Sts2 (Small tumor on seedlings 2). Expression of *Sts2* starts from 1 dpi (days post infection), and peaks between 2-4 dpi, when the tumor induction is initiated **(Supplementary Fig. 1b)** ^22^. Wheatgerm agglutinin-Alexa Fluor 488 (WGA-AF488) and propidium iodide co-staining showed no obvious colonization defect in the CR-Sts2 mutant compared to SG200 at 2 dpi **(Fig. 1b)**. We also monitored the growth of the mutant by relative fungal biomass quantification to 6 dpi **(Fig. 1c)**, the timepoint when leaf tumors get visible. Here, we did not detect a growth difference between CR-Sts2 and SG200, suggesting that the defect of tumor formation is not caused by a compromised biotrophic growth of the mutant, i.e. that Sts2 is probably not an inhibitor of host immunity.

**Fig1.**
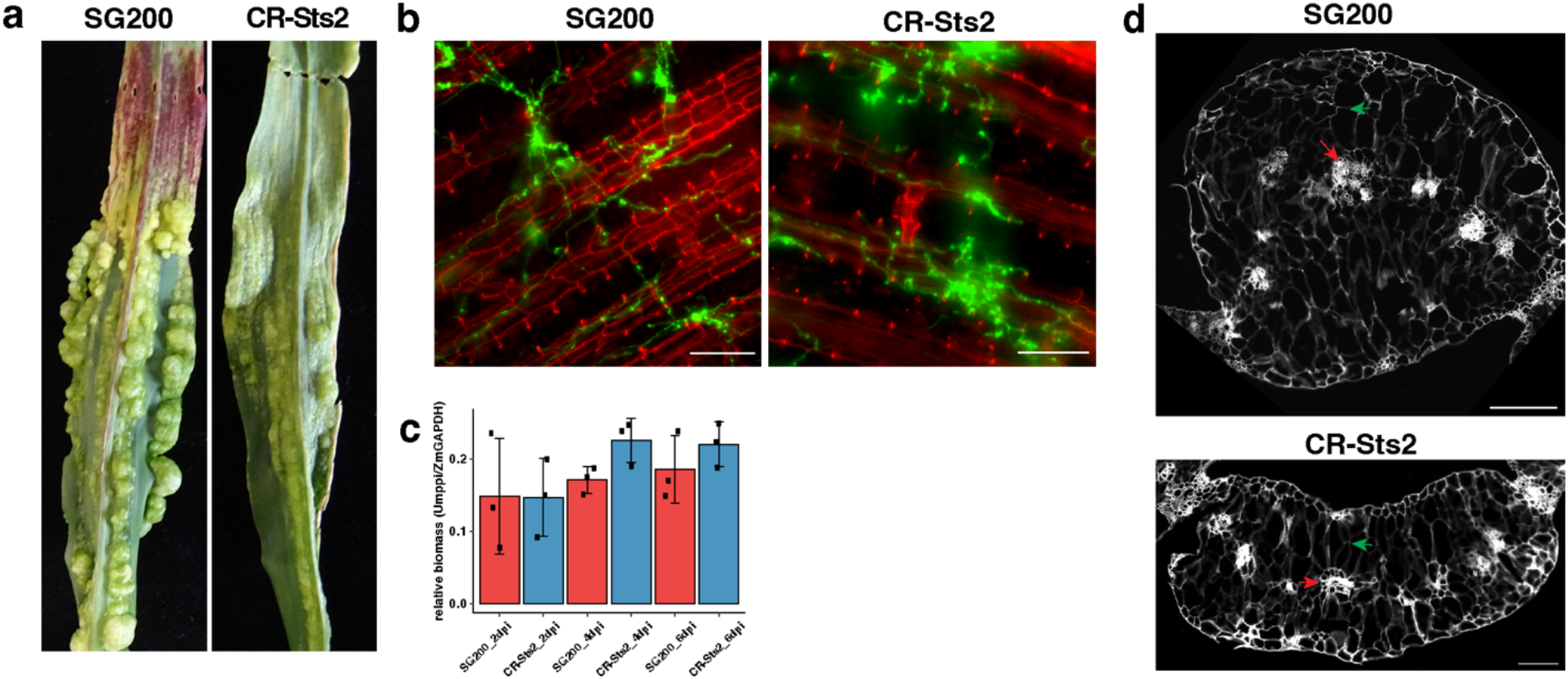
Sts2 regulates the hyperplastic tumor cell formation. **a,** Image shows the difference of the phenotype between SG200 and CR-Sts2 in maize leaf infections at 12 dpi, representative leaves were photographed. **b,** Microscopic images show the infected maize cells from SG200 and CR-Sts2 at 2 dpi. Green indicates hyphae from WGA-AF488 staining, and red is cell wall stained by propidium iodide. Representative images are shown. **c,** qPCR shows the relative biomass between SG200 and mutant. **d,** Transverse leaf sections from 12 dpi illustrate the different tumor cells in SG200 and CR-Sts2 infected leaves between two large lateral veins. The cell wall autofluorescence (DAPI channel) is shown. The red arrows point out the hyperplasia tumor cells at the original bundle sheath position. The green arrows denote the hypertrophy tumor cells. Scale bar = 200 μm.

To further investigate how tumor formation is affected in the CR-Sts2 mutant, we embedded 12 dpi leaves for transverse sectioning. In SG200 infected samples, clusters of small hyperplasia tumor cells from *de novo* division of bundle sheath cells can be observed. These were surrounded by 5-6 layers of supernumerary hypertrophic cells from enlarged divided cells and mesophyll **(Fig. 1d, Supplementary Fig. 1b**), which is in accordance to our previous observation^13^. Contrary, in CR-Sts2 infected plants, the *de novo* division was restricted, which resulted in depletion of hyperplasia cells, as well as reduced layers of hypertrophic cells **(Fig. 1d, Supplementary Fig. 1b)**. In brief, knock-out of Sts2 causes a premature stop of bundle sheath cell division, which blocks further tumor expansion.

### Sts2 is secreted and translocated into host nucleus

Sts2 encodes a small protein containing 183 amino acids (aa), including a 27 aa N-terminal signal peptide (SP). To test for secretion, we expressed Sts2-mcherry in the CR-Sts2 mutant under control of the *pit2* promoter, which confers a high expression level throughout plant infection **(Fig. 2a)**^25^. In confocal microscopy, Sts2-mcherry accumulated on the edge of biotrophic hyphae (similar to the effector control Pit2-mcherry^25^) indicating secretion during maize infection. **(Fig. 2a, b).** In contrast, the mcherry alone was localized in the cytoplasm of hyphae cell **(Fig. 2c)**. Next, we determined the subcellular localization of Sts2 in the plant by transient expression of Sts2^DSP^-GFP in *Nicotiana benthamiana* **(Fig. 2d)**. The specific nuclear localization of Sts2^DSP^ suggests that Sts2 might be translocated into the host cells upon secretion by *U. maydis* **(Fig. 2d)**. To test this, we complemented the *U. maydis* CR-Sts2 mutant with a Pro^Sts2^::Sts2-3×HA construct. The expression of the Sts2-3xHA fusion protein fully restored virulence, indicating that the C-terminal HA-tags did not affect the virulence function of Sts2 **(Fig. 3g)**. 4 dpi leaves from Pro^Sts2^::Sts2-3×HA infection were collected and followed up with cell fractionation to detect subcellular localization of Sts2-3×HA **(Fig. 2e)**. By western blot, we could detect Sts2 in the nuclear component, more precisely in the fraction associated with chromatin pellet after nucleoplasm extraction **(Fig. 2e)**. Together, this data shows that Sts2 is a secreted *U. maydis* effector which is translocated into the host nucleus.

**Fig2.**
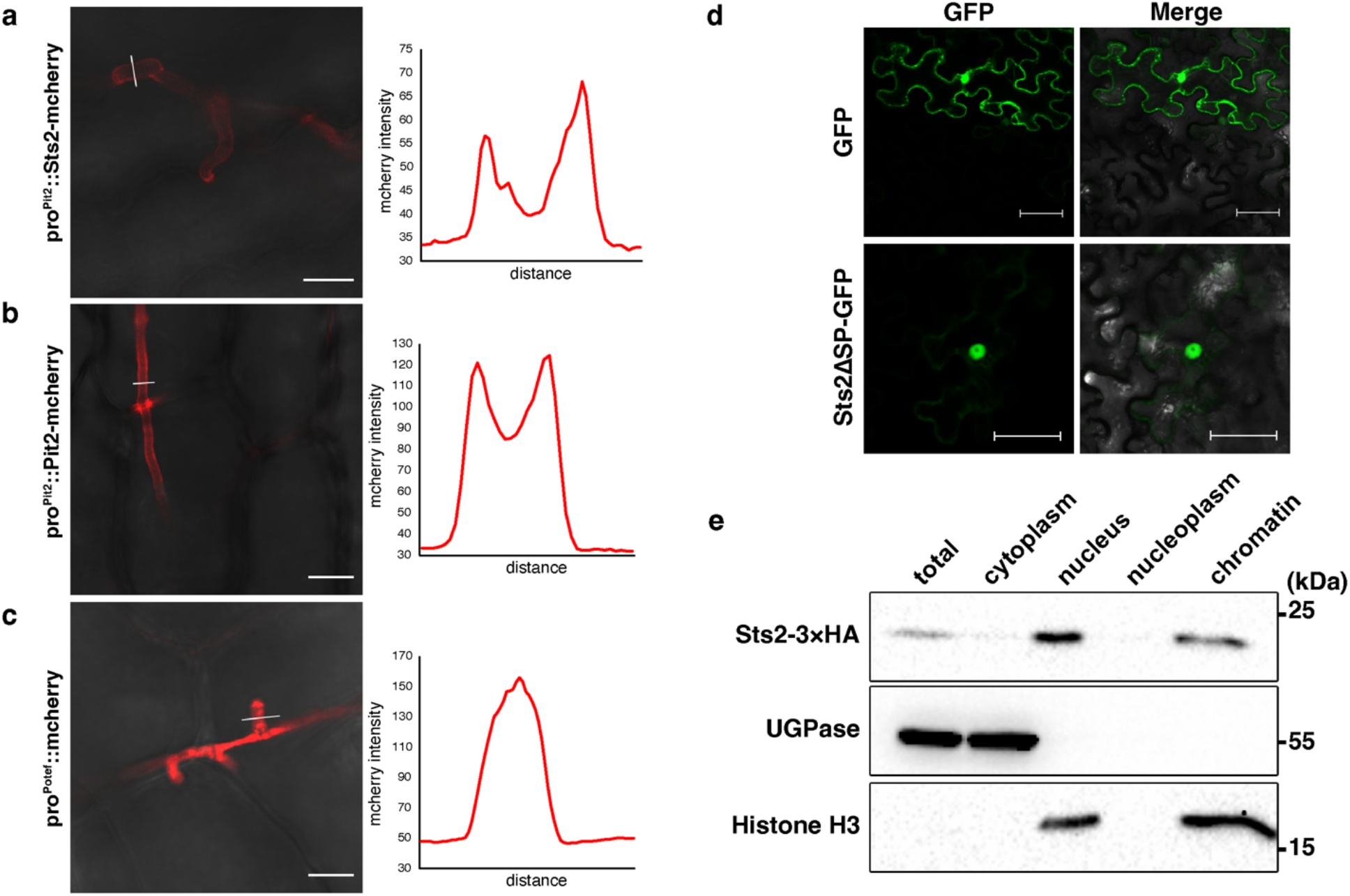
Sts2 is secreted and translocated into the host cell nucleus. **a-c,** *U. maydis* hyphae expressing Sts2-mcherry **(a)**, Pit2-mcherry **(b)** and **(c)** mcherry at 2 dpi. The plots are the mcherry intensities measured from the solid lines indicated in the photos. Scale bar = 10 μm. **d,** Localization of GFP and Sts2-GFP in *N. benthamiana.* Scale bar = 50 μm. **e,** Western blot detection of Sts2-3×HA in different cell fractions from 3 dpi maize leaves. UDP-Glucose-Pyrophosphorylase (UPase) and histone H3 were used as the markers of cytoplasmic and chromatin-associated fractions, respectively. The positions of molecular weight ladder are shown on the right.

**Fig3.**
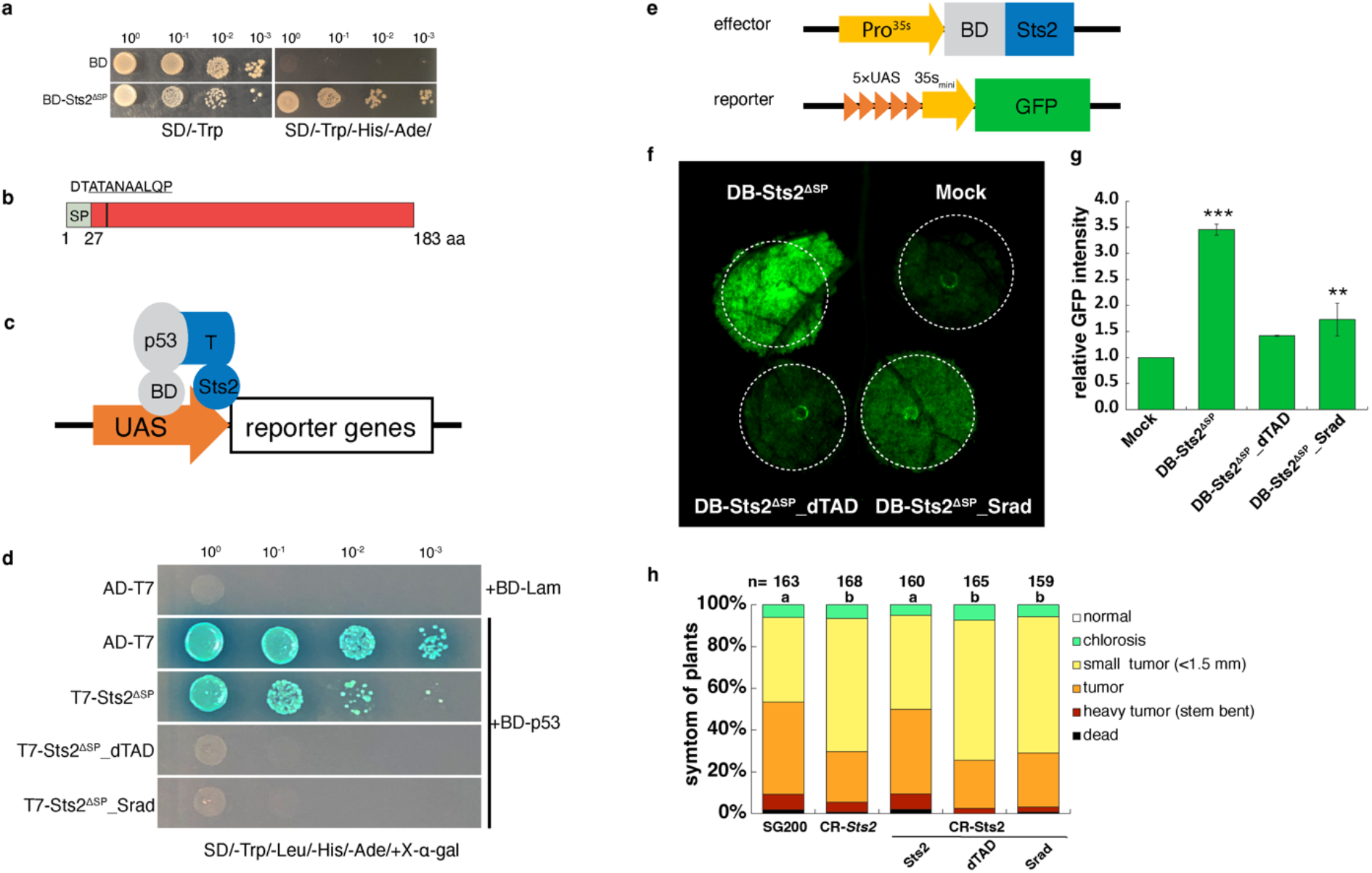
Sts2 is a transcriptional activator. **a,** The image shows the BD-Sts2^ΔSP^ transformed yeast grown on dropout SD medium plate. **b,** Domain arrangement of Sts2. The vertical line shows the position of overlapping TADs, and the amino acid sequences are shown above. The underlined sequences indicate the domain mutated or deleted in the following experiments. **c,** Diagram explains the co-transformation of BD-p53 and T7-Sts2^ΔSP^ and the interaction between p53 and T7 antigen brings Sts2^ΔSP^ to the promoter of reporter genes to activate their expressions. **d,** Image showing the growth of yeasts transformed with corresponding constructs on dropout SD plate. **e,** Diagram showing the construct used for agroinfiltration. **f,** GFP detection in *N. benthamiana* leaf co-infiltrated with the corresponding constructs together with Pro^5×UAS-35Smini^::GFP. The dashed circles indicate the infiltration area. **g,** Relative GFP intensity from three biological replications. The intensity was normalized to mock. Data shown are the mean value ±SD from 3 biological replication of 3 independent infiltrations. Student’s *t-test* was used for significance analysis. **, *p*<0.01. ***, *p*<0.001. **h,** The disease symptoms of SG200, CR-Sts2 and complementation strains with Sts2 and different Sts2 variants. N is the total number of plants infected from 4 independent infections. The letters are the significance levels between different strains. The Tukey Multiple Comparison test with Bonferroni *p* adjusted was performed to determine the significance.

### Sts2 has *trans* activator function

To elucidate the molecular function of Sts2, we attempted to use the yeast two hybrid system to identify potential maize interactors. When we fused Sts2 with Gal4 DNA binding domain (BD), the recombinant protein exhibited a strong activation of reporter genes and grew on dropout plate **(Fig. 3a)**, which implied that Sts2 may have a transactivation function. Using the 9aaTAD tool (https://www.med.muni.cz/9aaTAD/), we identified two overlapping nine aa (DTATANAAL and ATANAALQP) transactivation domains (TAD) in Sts2 **(Fig. 3b)**. To test the function of the predicted TAD, we mutated the domain ATANAALQP, since it was identified by “less stringent” and “pattern for clusters” models. We generated two different mutants by changing ATANAALQP into the corresponding aligned amino acids (EQAREHIQA) of the orthologous protein Sr10075 (Srad) from *S. reilianum*, or deleting the entire motif (dTAD). To minimize the noise expression of the reported genes in yeast, we adapted the positive interaction control p53 and T antigen used in yeast two hybrid assay by fusing Sts2 with T antigen to replace the Gla4 activation domain **(Fig. 3c)**. The interaction between p53 and antigen T brings Sts2 to the proximal of the promoter upstream the reporter genes thus activating their expression **(Fig. 3d)**. As expected, co-transformation of BD-p35 with T-Sts2^ΔSP^_Srad or T-Sts2^ΔSP^_dTAD failed to grow on dropout plate, indicating the loss of transactivation activity in the mutant proteins **(Fig. 3d)**. In a next step, we tested the *trans* activation of Sts2 *in planta.* To this end, we set-up an effector-GFP reporter system in *Nicotiana benthamiana* **(Fig. 3e)**. In consistency with the yeast experiments, DB-Sts2^ΔSP^ significantly induced GFP expression in *N. benthamiana* driven by the Pro^5×UAS-35Smini^ **(Fig. 3f).**This induction was significantly reduced or completely abolished, when the TAD was mutated or deleted, respectively **(Fig. 3g)**. Thus, Sts2 is a transcriptional activator *in-planta*, and this activity depends on its TAD motif. A crucial question is, if this activity is necessary for the virulence function of Sts2. To test this, we genetically complemented *U. maydis* CR-Sts2 with either, wild-type Sts2, Sts2_Srad or Sts2_dTAD mutant under its native promoter and used the resulting strains for plant infection assays. While wild-type Sts2 complementation completely restored virulence, strains expressing TAD-mutated versions showed reduced tumor formation similar to the CR-Sts2 **(Fig. 3h)**. This suggests that the *trans* activation function of Sts2, mediated by the TAD, is crucial for its virulence function.

### Sts2 interacts with ZmNECAP1, a putative maize transcriptional activator

Sts2 does neither contain canonical DNA binding domains, nor nuclear localization signals, which suggests that it is a transcriptional activator, and participates in the hosts gene regulation network. To understand how Sts2 activates host gene expression, we performed a co-immunoprecipitation (co-IP) to identify the interacting host proteins. For this, CR-Sts2 expressing the Sts2-3×HA was infected to maize leaves for subsequent sample preparation. As a control, *U. maydis* SG200 expressing GFP-3’HA (with the N-terminal secretion signal of Sts2, and expressed under control of the native sts2 promoter) was used. The immunoprecipitated proteins were subjected to mass spectrometry for protein identification. In total, 7 maize proteins were exclusively detected in Sts2-3’HA samples but not in the GFP-3’HA controls from at least 3 independent biological replicates **(Supplementary Table 1)**. Of these, we could independently confirm the interaction of Sts2 with Zm00001eb221890 both by co-IP **(Fig. 4a)** and split luciferase complementation assay in *N. benthamiana* **(Fig. 4b)**. Zm00001eb221890, which is predicted as an adaptin-ear-binding coat-associated protein 1 (ZmNECAP1 hereafter) containing a pleckstrin homology domain, shows expression in maize leaves, and displays significant transcriptional induction upon *U. maydis* infection **(Supplementary Fig. 2a).** Subcellular localization identifies ZmNECAP1 in the cytoplasm, nuclear membrane, and it co-localizes with Sts2 in the plant nucleus **(Fig. 4c, Supplementary Fig. 2b)**.

**Fig4.**
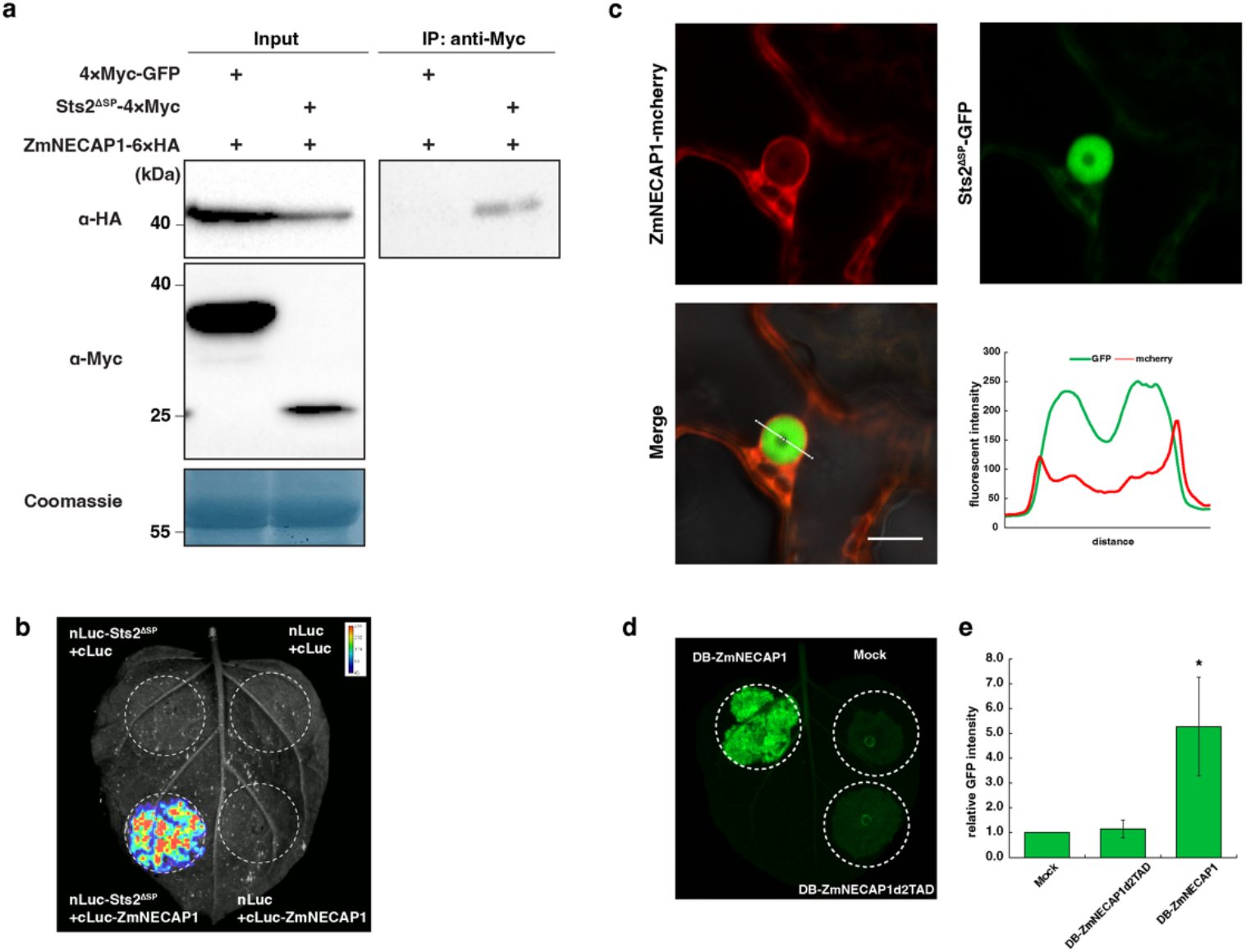
Sts2 interacts with a novel maize transcription activator ZmNECAP1. **a.** Western blot detection of co-immunoprecipitation experiment in *N. benthamiana.* **b.** Split luciferase complementary assay in *N. benthamiana* shows the interaction between Sts2^ΔSP^ and ZmNECAP1. The circles indicate the infiltration area. **c.** Microscope photos show the co-localization of Sts2^ΔSP^ and ZmNECAP1 in nucleus. The red is ZmNECAP1-mcherry, and the green is Sts2^ΔSP^-GFP. The GFP and mcherry intensities were measured from the solid white line as shown in the “Merge”. Scale bar = 10 μm. **d.** The image shows the expression of DB-ZmNECAP1 activated Pro^5×UAS-35Smini^::GFP. The dashed circles indicate the infiltration area. **e.** The bar plot shows the GFP intensity from three biological replications of three independent infiltrations. Data shown are the mean value ±SD and normalized to the Mock. Student’s *t-test* was used to determine the significance. “*”, *p*<0.05.

To our surprise, ZmNECAP1 contains two separated TADs **(Supplementary Fig. 2c)**. Similar to Sts2, a DB-ZmNECAP1 activated the reporter GFP expression under Pro^5×UAS-35Smini^ in *N. benthamiana* **(Fig. 4d, f),**while deletion of the TADs completely abolished this activity **(Fig. 4d, f)**. Thus, not only Sts2, but also its maize interactor ZmNECAP1 is a transcriptional activator.

### Sts2 activates maize leaf developmental regulators for tumor formation

To determine the host genes potentially regulated by Sts2, we conducted RNA-sequencing. For this purpose, *U. maydis* SG200, CR-Sts2 infected and mock-treated leaves from 3 and 6 dpi were sampled and analyzed. The infection of *U. maydis* dramatically altered maize leaf transcriptome. More than 45.2% of genes (11394 and 13593 genes at 3 and 6 dpi, respectively) were differentially expressed between mock treated and SG200 infected samples **(Supplementary Figure. 3a)**. In general, CR-Sts2 triggered a similar maize transcriptional change at different timepoints to that by SG200 **(Supplementary Figure. 3a)**, which confirms that mutation of Sts2 did neither affect the biotrophic growth of the mutant, nor trigger increased maize immune responses. In total, 5035 genes and 2370 genes were up-regulated or down-regulated, respectively, during the *U. maydis* infection, regardless of the genotype and time point **(Supplementary Figure. 3a, b)**. Gene ontology (GO) enrichment analysis identified genes involved in several cell cycles related biological processes were both up-regulated by SG200 and CR-Sts2 infection **(Supplemental Table 2)**, which is in line with that the *de novo* cell division of bundle sheath were initiated, but prematurely stopped by knockout as shown by transverse section microscopic photo.

To further elucidate key factors that control the sustained hyperplasia cell division, we compared the maize DEG between SG200 and CR-Sts2 infection samples in pairwise. In total, 435 and 465 genes were significantly up-regulated in SG200 samples at 3 and 6 dpi respectively, compared to 271 and 339 genes in CR-Sts2 samples **(Fig. 5a, b)**. GO enrichment analysis revealed that at 3 dpi, genes involved in the “stem cell population maintenance” and “meristem maintenance” are specifically activated by *U. maydis* infection in presence of Sts2 **(Fig. 5c)**. At 6 dpi, in addition to “meristem maintenance”, several developmental processes were up-regulated depending on Sts2 **(Fig. 5c)**. Interestingly, from the 83 genes that were consistently, significantly up-regulated in SG200 infected samples at both time points **(Fig. 5b)**, we identified the expression level of several transcription factors and activators were intensified in the presence of Sts2, including *ZmGRF3* (growth-regulating factor 3), *ZmGIF1* (growth-regulating-factor-interacting factor 1), *ZmYAB1* (YABBY1), *ZmSHR1* (short root 1), *ZmWOX5b* (WUSCHEL-Homeobox-transcription factor 5b) and *ZmANT1* (AINTEGUMENTA 1) **(Fig. 5d).**These genes are highly expressed in the dividing zone during leaf development^26^ and in maize embryonic leaf cells^27,28^. ZmGRF interacts with ZmGIF, together to determine the shoot meristem in maize^29–31^, where ZmANT1, ZmSHR1 and ZmYAB1 are regulators of Kranz anatomy development^27,28,32^ and WOX members are well known highly expressing in embryo like cells^33^. Overexpression of some of these genes alone can lead to excess cell division in Arabidopsis^34^, rice^35^ and Medicago^36^. While these genes are highly induced upon SG200 infection compared to mock samples, knock out of Sts2 resulted in a strongly reduced expression **(Supplementary. 4)**. Furthermore, CR-Sts2 strains complemented with mutated TAD (Srad and dTAD) also failed to activate the expression of these genes to the levels observed for SG200. Similarly, *U. maydis* strains where UmSts2 was replaced by the open reading frames of the two *S. reilianum* Sts2 orthologs (KI_sr10075 and KI_sr10079) could not induce these maize genes during infection. **(Fig. 5e)**. Together, this shows that the activation of these maize genes requires Sts2 with its functional transactivation motif.

**Fig5.**
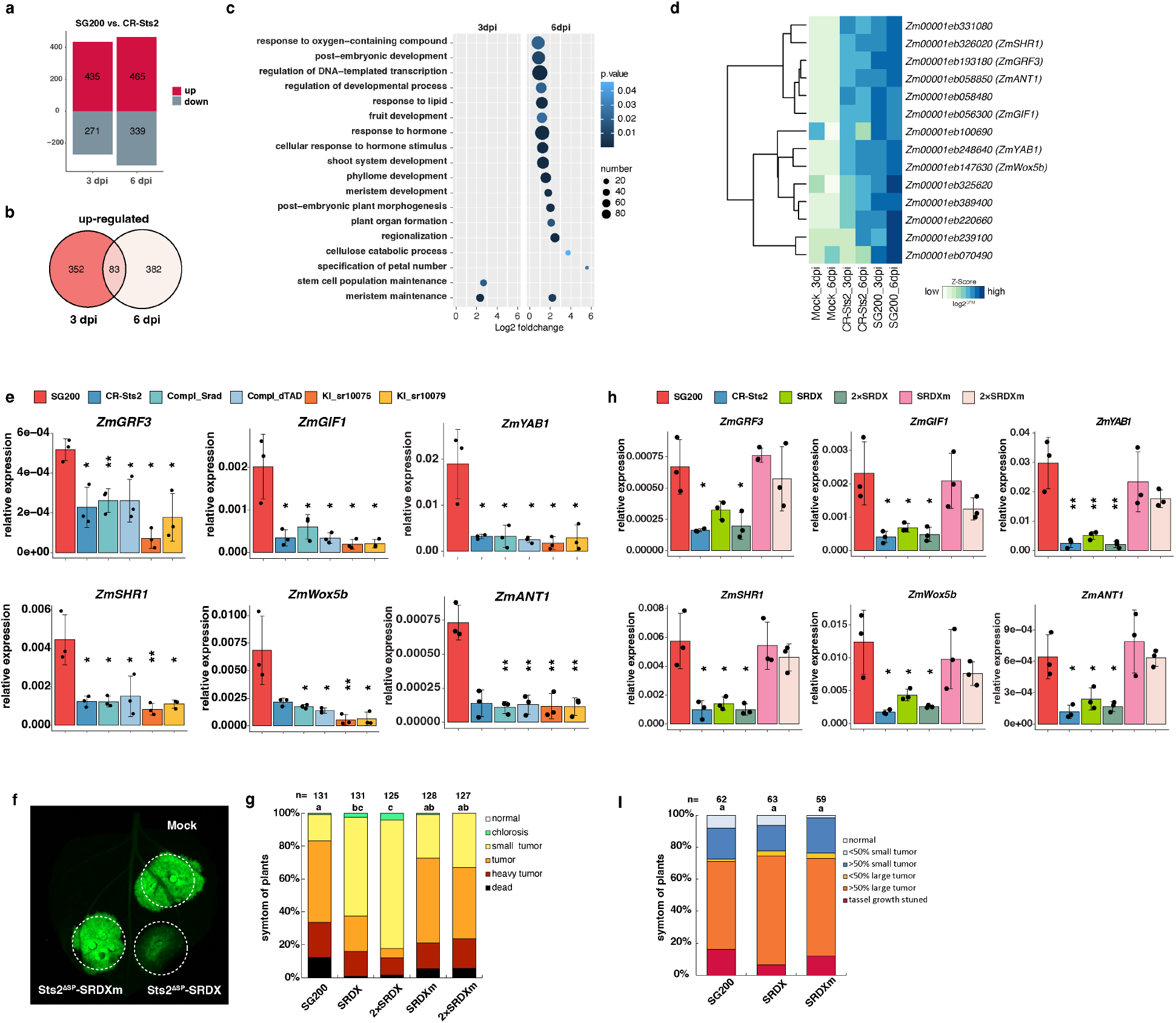
Sts2 activates regulators of maize leaf development. **a,** Bar plot shows number of genes significantly up- or down-regulated in SG200 compared to CR-Sts2 at 3 and 6 dpi. **b,** Venn diagram shows a total of 83 genes were consistently up-regulated upon SG200 infection from both timepoints compared to CR-Sts2. **c,** The dot plot shows GO enrichment analysis of DEGs. **d,** The heatmap shows the expression levels of differentially expressed transcription regulators at both timepoints between SG200 and CR-Sts2 infection. The Z score normalization was performed on the log2CPM (counts per million) of each gene across samples. **e,** The quantitative RT-PCR results show the expression levels of *ZmGRF3*, *ZmGIF1*, *ZmYAB1*, *ZmSHR1*, *ZmWox5b* and *ZmANT1* upon infection of different strains. Data shown are the mean value ±SD from 3 biological replications of 3 independent infections, the solid points indicate the value of each replication. Student’s *t-test* was used for significance test of the expression levels between SG200 and mutants. “*”, *p*<0.05, “**”, *p*<0.01. **f,** Sts2^ΔSP^-SRDX suppresses the transactivation function of ZmNECAP1. A level2 MoClo construct containing the Pro^5×UAS-35Smini^::GFP and DB-ZmNECAP1 was co-infiltrated with respective construct shown in the photo. The dashed circles indicate the infiltration area. **g,** The disease scoring of GB infected by SG200, and SG200 overexpressing Sts2-SRDX and Sts2-SRDXm. The experiment was repeated three times with independent infection, n is the total number of infected plants. The letter denotes the significance level between each strain. The Tukey Multiple Comparison test with Bonferroni *p* adjust was performed to determine the significance. **h**, qPCR quantification of the expression levels of leaf developmental regulators upon SG200, SG200 overexpression Sts2-SRDX and Sts2-SRDXm infection at 6 dpi. Data shown are the mean value ±SD from 3 biological replications of 3 independent infections, the solid points indicate the value of each replication. Student’s *t-test* was used for significance test of the expression levels between SG200 and mutants. “*”, *p*<0.05, *p*<0.01.

To test, if the reduced expression of these genes is specific to Sts2 regulation rather than being a consequence of compromised tumor formation, we checked the expression of these Sts2-induced genes in other *U. maydis* effector mutants being compromised in tumorigenesis. Infection of maize inbred line CML322 with an *U. maydis* deletion mutant for the effector gene *UMAG_02297* resulted in reduced tumors^37^. Re-analysis of corresponding RNA-seq data reveals that *ZmGIF1*, *ZmGRF3*, *ZmYAB1*, *ZmSHR1*, and *ZmANT1* were not significantly changed in the *UMAG_02297* knockout compared to SG200, although expression of *ZmWOX5b* could not be detected **(Supplementary Fig. 5a)**. More importantly, qPCR showed that induction of the Sts2-induced maize genes is not affected upon *U. maydis* ΔSee1 mutant infection (**Supplementary. 5b**), which displays a similar arrested hyperplasia tumor division phenotype to CR-Sts2^13,20^. All together, these findings confirm a specific induction of the identified maize genes by Sts2.

Following our conclusion that Sts2 induces the expression of maize genes required for tumorigenesis, we hypothesized that the transcriptional repression of Sts2-regulated maize genes would result in an inhibition of tumor formation. However, as the maize mutants of these genes are not available, we decided for the alternative approach to fuse a transcriptional suppressor SRDX motif with Sts2. The SRDX motif was shown to turn transcriptional activators into suppressors and moreover, it could inhibit the *trans* activation of its interacting transcription factors /activators^38,39^. Thus, an Sts2-SRDX fusion is expected to act as a dominant suppressive effector. Indeed, in the effector-GFP reporter system, DB-ZmNECAP1 activated the expression of GFP under the control of Pro^5×UAS-35Smini^::GFP as shown above **(Fig. 4d,e)**. However, upon co-infiltration with Sts2^ΔSP^-SRDX, this activation was suppressed by the interaction between ZmNECP1 and Sts2^ΔSP^-SRDX as expected, but not by the Sts2 fused with mutated SRDX motif (SRDXm) **(Fig. 5f)**. To test the effect of Sts2-SRDX mediated repression in the actual *U. maydis* - maize interaction, we generated *U. maydis* strains over-expressing Sts2-SRDX (or Sts2-SRDXm) to outcompete the native Sts2 and infected them to maize seedlings to evaluate the tumor formation. At 12 dpi, *U. maydis* strains expressing Sts2-SRDX, but not Sts2-SRDXm, showed significantly reduced the tumor formation. This reduction seemed to be dose dependent, as multiple integration of the Sts2-SRDX overexpression construct had an additive effect, resulting in further decreased tumor formation **(Fig. 5g)**. qPCR showed that the transcript levels of Sts2-dependent leaf developmental regulators were significantly inhibited by Sts2-SRDX. Contrary, Sts2-SRDXm did not significantly affect expression of the maize genes, independent of its copy number **(Fig. 5h)**. Notably, overexpression of Sts2-SRDX or Sts2-SRDXm did not completely suppress or further increase the expression of these genes **(Fig. 5h).**

We found previously that Sts2 contributes to fungal virulence during leaf infection, but not on tassel, where *U. maydis*-induced tumor formation results from redirecting the intrinsic cell proliferation without an oncogenic activity, as it is seen in leaf tumors ^13,24^. In line with this, tassel infection of the Sts2-SRDX and Sts2-SRDXm mutants showed no reduction in virulence compared to strain SG200 **(Fig. 5i).**This underpins that the restricted tumor formation caused by Sts2-SRDX is not consequence of a general inhibition of *U. maydis* virulence, but results from the suppression of tissue specific host developmental regulators. Taken together, we conclude that the Sts2 effector is translocated from fungal hyphae to plant nuclei, where it contributes to tumor formation by activating the expression of leaf development regulators.

## Discussion

*U. maydis* induced tumor formation is a complex and still poorly understood process. During this process, *U. maydis* secretes a group of the tumorigenic effectors which not only prime highly differentiated bundle sheath cells for *de novo* cell division, but also sustains such division to increase the tumor size. Until now, See1 is the only effector which has been shown to re-activate the DNA synthesis, a prerequisite of cell division^20^. Surprisingly, *S. reilianum* SrSee1 protein resembles this function in *U. maydis*, but not UhSee1 from the barley smut fungi *Ustilago hordei*^20,40^. Here, we disclose Sts2 as a tumorigenic effector and, to our best knowledge, the first functionally validated fungal effector which acts as a transcriptional activator in the host. Moreover, we found that the virulence function of Sts2 depends on its transcriptional activation function. Compared to TALE effectors of the bacterial pathogen *Xanthomonas sp*^3^, Sts2 lacks a repetitive DNA binding domain, which may be related to the generally small size of fungal effectors to facilitate the secretion into host cell via conventional secretory pathway. The lack of a DNA binding domain also implies that Sts2 needs to recruit host components to form a transcriptional complex to activate target gene expression. We found that Sts2 interacts with ZmNECAP1, a functionally uncharacterized plant transcriptional activator. Here, future investigations on the function of ZmNECAP1 during *U. maydis*-induced tumor formation and leaf development will aim to elucidate whether it is a co-activator with Sts2, or rather a scaffold required by Sts2 to target a DNA binding protein are needed.

*U. maydis* infection comprehensively reprograms the leaf developmental process to form tumors, which makes it hard to pinpoint the key host factors related to tumorigenesis. In this study, we show that Sts2 amplifies the expression of a group of plant transcription factors and activators regulating leaf development, after the developed bundle sheath cells are primed for division. We hypothesize that the Sts2-regulated genes are the potential executors to maintain hyperplasia tumor propagation. Accordingly, Sts2-SRDX specifically suppresses these genes, leading to compromised tumor formation on maize leaves, but not on tassel. It will be intriguing to know whether these genes are directly regulated by Sts2, or whether their regulations are decided downstream of Sts2 by yet unknown transcription factors. The results presented in this study demonstrate the potential of microbial effectors as molecular tools to help us explore complex cellular processes in host organisms, such as the network of co-regulation of leaf development regulators. Above all, we continue to be fascinated by the power of evolution to produce the molecular diversity of pathogen-host interactions necessary to adjust highly complex processes such as tumor formation in a cell type-specific manner.

## Materials and Methods

### Strains and plant material, growth conditions

All mutants were generated in *U. maydis* solopathogenic strain SG200 in this study. The *U. maydis* strains were grown on PD-agar plate at 28°C or YEPS light liquid medium at 28°C, 200 rpm. The maize cultivar Golden Bantam was used for infection (otherwise indicated). The plants were grown at controlled greenhouse or phytochamber with 16 hr light at 28°C and 8 hr dark at 22°C. The CRISPR mutagenesis of Sts2 was done as previous described^23^ by the oligo listed in **Supplementary Table 3**. The resulting strains were sent for sequencing to confirm the open reading frame shift and premature stop. For Sts2-SRDX or SRDXm, the amino acid sequence of SRDX (LDLDLELRLGFA) ^39^ and SRDXm (FDPDQEARFGFA)^38^ were first codon optimized in Eurofin online tool, and then assembled with Sts2 open reading frame and Pit2 promoter by Gibson assembly. The constructs were then introduced in SG200 by *ip* integration and further analyzed by southern blot and qPCR to check the integration and copy number (data not shown). The complementary of all CR-Sts2 with Sts2 and Sts2 mutants were done in a similar method.

### Transactivation activity test

For the autoactivation test in yeast, the yeast strain AH109 was used. The StSts2_28-183_ was amplified and cloned into pGBKT7 to fuse with Gal4 BD domain, or into pGADT7-T to replace Gal4 AD domain and in open reading frame with antigen T protein. The resulting plasmids were transformed into AH109 alone or with pGBKT7-p53, respectively. The transformation and drop assay were done according to the protocol from manufacturer.

For the autoactivation test in *Nicotiana Benthamiana*, the Sts2_28-183_, Gal4-BD domain and promoter 5×UAS-35Smini were cloned into MoClo system with BsaI and BpiI domestication^41^, and further used for modular cloning of the corresponding. The constructs were transformed into agrobacterium strain GV3101 by electroporation. The resulting agrobacterium was grown overnight until OD_600_ between 1-2, and suspended in infiltration buffer (10 mM MgCl_2_, 10 mM MES pH5.6, 200 μM acetosyringone) to OD_600_=3, incubated in dark for 2 hr. Before infiltration, an equal volume of each strain was mixed with p19 strain to final OD=1. Leaves from 3 dai (days after infiltration) were used for GFP detection by ChecmiDoc MP machine (BioRad), and ImageJ Fiji was used for quantification of GFP intensity. The GFP intensity was normalized to the mock infiltration.

### Maize infection and disease scoring

For seedling infection, seven-days-old Early Golden Bantam (EGB) seedlings or six-days-old Golden Bantam (GB) seedlings were used. The infection and disease scoring were done as previously described^23^. The disease index was used for statistic test, and Tukey multiple comparison test with Bonferroni adjustment was used for significance test. For tassel infection and scoring were done on maze cultivar Gaspe Flint according to Redkar *et. al*.^42^. For significance test, a similar disease index was used as in seedling infection by assigning index 9 to tassel growth stunted, >50% large tumor (7), >50% small tumor (5), <50% large tumor (3), <50% small tumor and normal tassel as 0. Then the disease index was used for statistic test, and Tukey multiple comparison test with Bonferroni adjustment was used for significance test. The biological replications were from independent infection experiments.

### Leaf staining, tissue embedding, sectioning and microscope

Wheat germ agglutinin-Alexa Fluor 488 (WGA-AF488) and propidium iodide co-staining was done according to previous description. For tissue embedding, around 1.8-cm leaf sections 3 cm below the infection sites were collected and embedded according to Alexandra M, 2017^13^. Leaf tissues were arranged in Peel-A-Way^™^ Einbett-Formen mold, and sectioned into 15 μm for microscope. The microscopy of staining tissues and transverse sections were done by using a Nikon Eclipse Ti inverted microscope with the Nikon Instruments NIS-ELEMENTS software. The GFP and mcherry microscopy were done by using a Leica TCS SP8 confocal laser scanning microscope.

### Co-immunoprecipitation and mass spectrometry in maize

Inoculums with OD_600_=3 and 0.1% tween-20 were used to infect EGB. At 3 dpi, 4 cm length of leaf sections 1 cm below the infection site were collected and ground into fine powder with liquid nitrogen. The powders were incubated in extraction buffer (50mM Tris-HCl pH7.5, 150mM NaCl, 10% glycerol, 2mM EDTA, 5mM DTT, 1% Triton X-100 and protease inhibitor) for 30 min on ice and centrifuged twice at 16,000 g, 4 °C for 30 min. 10 μl of anti-HA magnetic beads (Pierce) were added into each supernatant and followed by 1 hr incubation at 4 °C with end-to-end rotation. Afterward, the beads were washed three times with extraction buffer and three times with extraction buffer without Triton X-100. In total, four replications from 4 independent infections were prepared and to MS analysis.

For MS analysis, dry beads were re-dissolved in 25 μL digestion buffer 1 (50 mM Tris, pH 7.5, 2M urea, 1mM DTT, 5 ng/μL trypsin) and incubated for 30 min at 30 °C in a Thermomixer with 400 rpm. Next, beads were pelleted and the supernatant was transferred to a fresh tube. Digestion buffer 2 (50 mM Tris, pH 7.5, 2M urea, 5 mM CAA) was added to the beads, after mixing the beads were pelleted, the supernatant was collected and combined with the previous one. The combined supernatants were then incubated o/n at 32 °C in a Thermomixer with 400 rpm; samples were protected from light during incubation. The digestion was stopped by adding 1 μL TFA and samples were desalted with C18 Empore disk membranes according to the StageTip protocol^43^.

Dried peptides were re-dissolved in 2% ACN, 0.1% TFA (10 μL) for analysis and diluted 1:10 for measurement. Samples were analyzed using an EASY-nLC 1000 (Thermo Fisher) coupled to a Q Exactive mass spectrometer (Thermo Fisher). Peptides were separated on 16 cm frit-less silica emitters (New Objective, 75 μm inner diameter), packed in-house with reversed-phase ReproSil-Pur C18 AQ 1.9 μm resin (Dr. Maisch). Peptides were loaded on the column and eluted for 115 min using a segmented linear gradient of 5% to 95% solvent B (0 min : 5%B; 0-5 min -> 5%B; 5-65 min -> 20%B; 65-90 min ->35%B; 90-100 min -> 55%; 100-105 min ->95%, 105-115 min ->95%) (solvent A 0% ACN, 0.1% FA; solvent B 80% ACN, 0.1%FA) at a flow rate of 300 nL/min. Mass spectra were acquired in data-dependent acquisition mode with a TOP15 method. MS spectra were acquired in the Orbitrap analyzer with a mass range of 300–1750 m/z at a resolution of 70,000 FWHM and a target value of 3×10^6^ ions. Precursors were selected with an isolation window of 1.3 m/z. HCD fragmentation was performed at a normalized collision energy of 25. MS/MS spectra were acquired with a target value of 10^5^ ions at a resolution of 17,500 FWHM, a maximum injection time (max.) of 55 ms and a fixed first mass of m/z 100. Peptides with a charge of +1, greater than 6, or with unassigned charge state were excluded from fragmentation for MS^2^, dynamic exclusion for 30s prevented repeated selection of precursors.

### Co-immunoprecipitation in *N. benthamiana*, subcellular fractionation, western blot and split Luciferase complementary assay

The agroinfiltration and the co-IP was done as described above. Leaf samples from 2 dai were collected and ground to fine powder. The protein was extract by the following buffer (50mM Tris-HCl pH7.5, 150mM NaCl, 10% glycerol, 10mM ETDA, 1 mM DTT, 1 mM PMSF, 1% IGEPAL CA-630 and protease inhibitor). The Myc-Trap magnetic agarose beads (ChromoTek) were used for immunoprecipitation.

The subcellular fractionation was done according to the method from Haring M, 2007^44^. Afterwards, the resulting nucleus pellet was split into two halves. One half was used for nuclei lysis followed as Chang L, 2018^45^ with some modification. The nuclei pellet was suspended in 100 μl glycerol buffer (20 mM Tris-HCl, pH 7.9, 50% glycerol, 75 mM NaCl, 0.5 mM EDTA, 1 mM DTT, 1 mM PMSF and Roche protease inhibitor cocktail), then 100 μl prechilled nuclei lysis buffer (10 mM HEPES, pH 7.6, 1 mM DTT, 7.5 mM MgCl_2_, 0.2 mM EDTA, 0.3 M NaCl, 1 M Urea, 1% IGEPAL CA-630, 1 mM PMSF, and Roche protease inhibitor cocktail) was added, vortexed and incubated on ice for 5 min. After centrifugation at 4°C with maximum speed, the supernatant was taken as nucleoplasm fraction and the pellet was washed twice with cold PBS buffer containing 1 mM EDTA. The anti-Myc and anti-HA antibody were used for detection by using ChecmiDoc MP machine (BioRad).

Different agrobacterium strains were mix with p19 to the final OD_600_=1, the leaves from *Nicotiana Benthamiana* were shortly rinsed in water, sprayed with 1 mM D-luciferin (Promega) and kept in dark for 10 min before detection by using ChecmiDoc MP machine (BioRad).

### DNA and RNA preparation and quantitative PCR

For RNA-seq, at least 15 leaves from individual plants were mixed as one sample, for biomass and gene expression quantification, one sample was mixed from at least 5 individual leaves. Each experiment was repeated three times from three independent infections. DNA was prepared by Buffer A (0.1 M Tris-HCl, 0.05 M EDTA, 0.5 M NaCl, 1.5% SDS) and purified by MasterPure Complete DNA and RNA Purification Kit Bulk Reagents (Epicentre, Madison, WI, USA). RNA was prepared by TRizol according to the manufacturer’s protocol, followed by DNaseI digestion. The cDNA synthesis was done by using RevertAid First Strand cDNA Synthesis Kit (Thermo Scientific). The qPCR was done by using GoTaq qPCR mix (Promega) and performed on CFX96 Real-Time PCR Detection System (Bio-Rad). 2^-ΔCt^ (Ct^UmPpi^-CT^ZmGAPDH^) and 2^-ΔCt^ (Ct^GOI^-CT^reference^) was used to determine the relative biomass and gene expression, respectively and Student’s *t-test* was used for statistical analysis of significance.

### MS and RNA-seq data analysis

For MS data analysis, the raw data were processed using MaxQuant software (version 1.6.3.4, http://www.maxquant.org/)^46^ with label-free quantification (LFQ) and iBAQ enabled^47^. MS/MS spectra were searched by the Andromeda search engine against a combined database containing the sequences from *Z. mays* (Zmays_284_Ensembl-18_2010-01-MaizeSequence.protein_primaryTranscriptOnly.fasta), the bait protein and sequences of 248 common contaminant proteins and decoy sequences. Trypsin specificity was required and a maximum of two missed cleavages allowed. Minimal peptide length was set to seven amino acids. Carbamidomethylation of cysteine residues was set as fixed, oxidation of methionine and protein N-terminal acetylation as variable modifications. Peptide-spectrum-matches and proteins were retained if they were below a false discovery rate of 1%. Statistical analysis of the MaxLFQ values was carried out using Perseus (version 1.5.8.5, http://www.maxquant.org/). Quantified proteins were filtered for reverse hits and hits “identified by site” and MaxLFQ values were log2 transformed. After grouping samples by condition only those proteins were retained for the subsequent analysis that had two valid values in one of the conditions. Two-sample *t*-tests were performed using a permutation-based FDR of 5%. The Perseus output was exported and further processed using Excel.

The RNA-seq was done in Novogene. Reads were filtered using the Trinity software (v2.9.1) option trimmomatic under the standard settings^48^ and then mapped to the reference genome using Bowtie 2 (v2.3.5.1) with the first 15 nucleotides on the 5’-end of the reads being trimmed^49^. The reference genome was the genome assembly of *U. maydis^50^* combined with the assembly of *Z. mays* B73 version 5^51^. Reads were counted using the R package Rsubread (v1.34.7)^52^. The edgeR package v.3.26.8 was used for statistical analysis and determine the differential gene expression by using the pairwise comparison generalized linear models (GLMs). Genes with log2 fold change>1 or <-1 and *p*<0.05 were considered as differentially regulated between SG200 and ΔSts2 samples. The differentially regulated genes between treatment were subjected to gene ontology analysis by using PLAZA 5.0 with default setting.

## Supporting information

Supplementary Figures with legends

## Data availability

All data that support the findings of this study which are not directly available within the paper (and its supplementary information files) will be available from the corresponding authors (GD, WZ) upon reasonable request. RNAseq raw data are publicly accessible in the NCBI Gene Expression Omnibus with accession number GSE225929. The mass spectrometry proteomics data have been deposited to the ProteomeXchange Consortium via the PRIDE^53^ partner repository with the dataset identifier PXD040350

## Acknowledgements

This project has received funding from the European Research Council (ERC) under the European Union’s Horizon 2020 research and innovation programme (grant agreement No 771035), as well as funding by the Deutsche Forschungsgemeinschaft (DFG, German Research Foundation) under Germany’s Excellence Strategy-EXC-2048/1-Project ID: 390686111 and Research Grant DFG-Az: DO 1421/3-3.

## Author contributions

WZ and GD designed the research; WZ performed all molecular experiment, virulence assay and RNA-seq data analysis; JRLD mapped the RNA-seq data; SCS and HN conduced MS and MS data analysis. WZ and GD wrote the paper with contributions from other authors.

## Supplementary Information

**Supplementary Fig1. Sts2 is induced during *U. maydis* infection and regulates hyperplasia tumor formation. a,** The disease symptoms of SG200, CR-Sts2 on maize cultivar Early Golden Bantam. N is the total number of plants infected from 3 independent infections. “*”, *p*<0.05. The Student’s *t-test* was used for statistic test. **b,** The expression of Sts2 in *U. maydis* FB1×FB2 during the whole biotrophic infection from published data. **c,** More microscope photos of transverse section from SG200 and CR-Sts2 at 12 dpi on maize cultivar Golden Bantam. Each photo represents the typical phenotype from individual plant.

**Supplementary Fig2. ZmNECAP1 is a plant transcriptional activator induced during *U. maydis* infection. a,** The expression of ZmNECAP1 is induced during *U. maydis* FB1×FB2 infection from previously published data. **b,** The subcellular localization of ZmNECAP1 in *N. benthamiana.* **c,** The domain arrangement of ZmNECAP1. Two vertical lines indicate the position of the separated two TADs and the amnio acid sequences are shown above.

**Supplementary Fig3. Differentially expressed maize genes between *U. maydis* SG200 vs mock. a,** The Venn diagrams show the numbers of differentially expression genes between SG200vs. Mock and CR-Sts2 vs. Mock at 3 and 6 dpi, respectively. **b,** The intersection diagrams show the gene numbers in different paired comparison.

**Supplementary Fig4. The expression of maize genes upon *U. maydis* infection in RNA-seq samples by qPCR.** The expression level of leaf developmental regulators detected by qPCR in RNA-seq samples. The value of 2^-ΔCt^ between gene of interest (GOI) and *ZmGAPDH* are calculated and plotted.

**Supplementary Fig5. The expression of maize genes upon different *U. maydis* effector mutants’ infection. a,** The relative expression levels of maize leaf developmental regulators upon SG200 and KO_UMAG_02297 infection in maize line CML322 at 3 dpi from published data^37^. The CPM (counts per million) are plotted. **b,** The expression of leaf developmental regulators in SG200 and ΔSee1 mutant infected Golden Bantam at 6dpi by qPCR. The values of 2^-ΔCt^ between GOI and *ZmGAPDH* are calculated and plotted.

**Supplementary Table 1. Mass Spectrometry data**

**Supplementary Table 2. Gene ontology analysis of up- and down-regulated genes during *U. maydis* infection**

**Supplementary Table 3. Oligos sequencing used in the study**

