## Supplementary Figures with legends for "A transcriptional activator effector of *Ustilago maydis* regulates hyperplasia in maize during pathogen-induced tumor formation"

### Figure S1

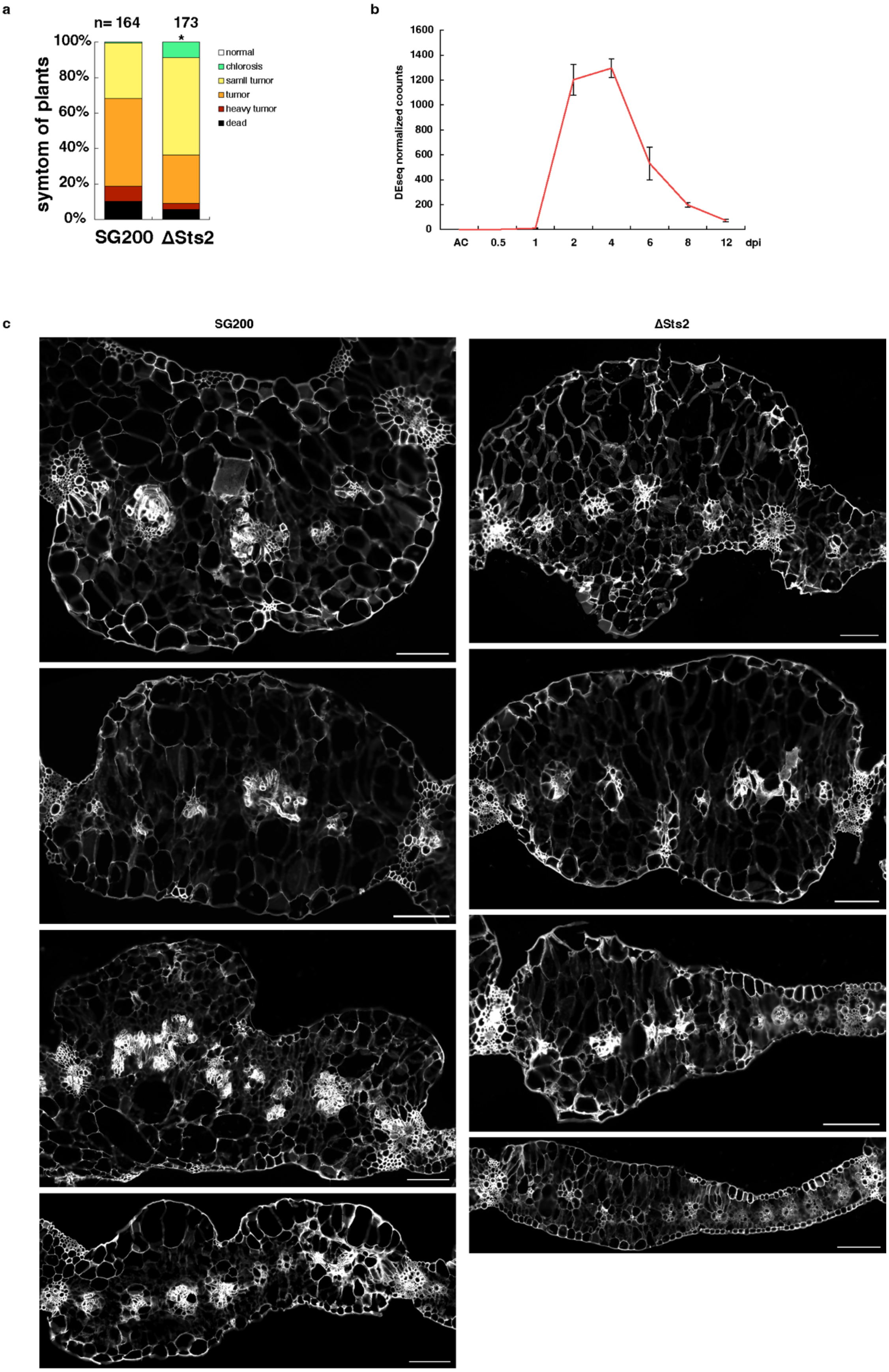
a
n= 164
173
100%
80%
60%
40%
20%
0%
symtom of plants
normal
chlorosis
samll tumor
tumor
heavy tumor
dead
SG200
ΔSts2
b
DEseq normalized counts
1600
1400
1200
1000
800
600
400
200
0
AC
0.5
1
2
4
6
8
12
dpi
c
SG200
ΔSts2

**Supplementary Fig1. Sts2 is induced during *U. maydis* infection and regulates hyperplasia tumor formation. a,** The disease symptoms of SG200, CR-Sts2 on maize cultivar Early Golden Bantam. N is the total number of plants infected from 3 independent infections. "*", $p<0.05$. The Student's *t-test* was used for statistic test. **b,** The expression of Sts2 in *U. maydis* FB1×FB2 during the whole biotrophic infection from published data. **c,** More microscope photos of transverse section from SG200 and CR-Sts2 at 12 dpi on maize cultivar Golden Bantam. Each photo represents the typical phenotype from individual plant.

### Figure S2

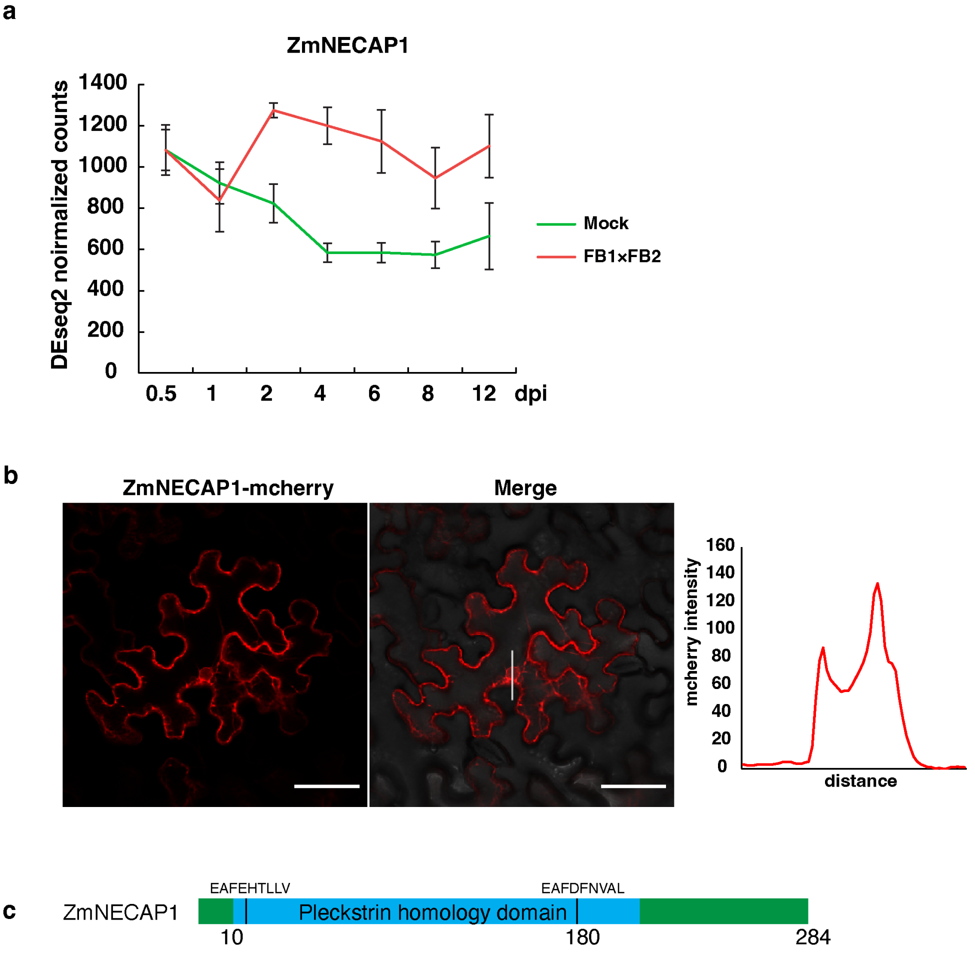


**Supplementary Fig2. ZmNECAP1 is a plant transcriptional activator induced during *U. maydis* infection. a,** The expression of ZmNECAP1 is induced during *U. maydis* FB1×FB2 infection from previously published data. **b,** The subcellular localization of ZmNECAP1 in *N. benthamiana.* **c,** The domain arrangement of ZmNECAP1. Two vertical lines indicate the position of the separated two TADs and the amnio acid sequences are shown above.

### Figure S3

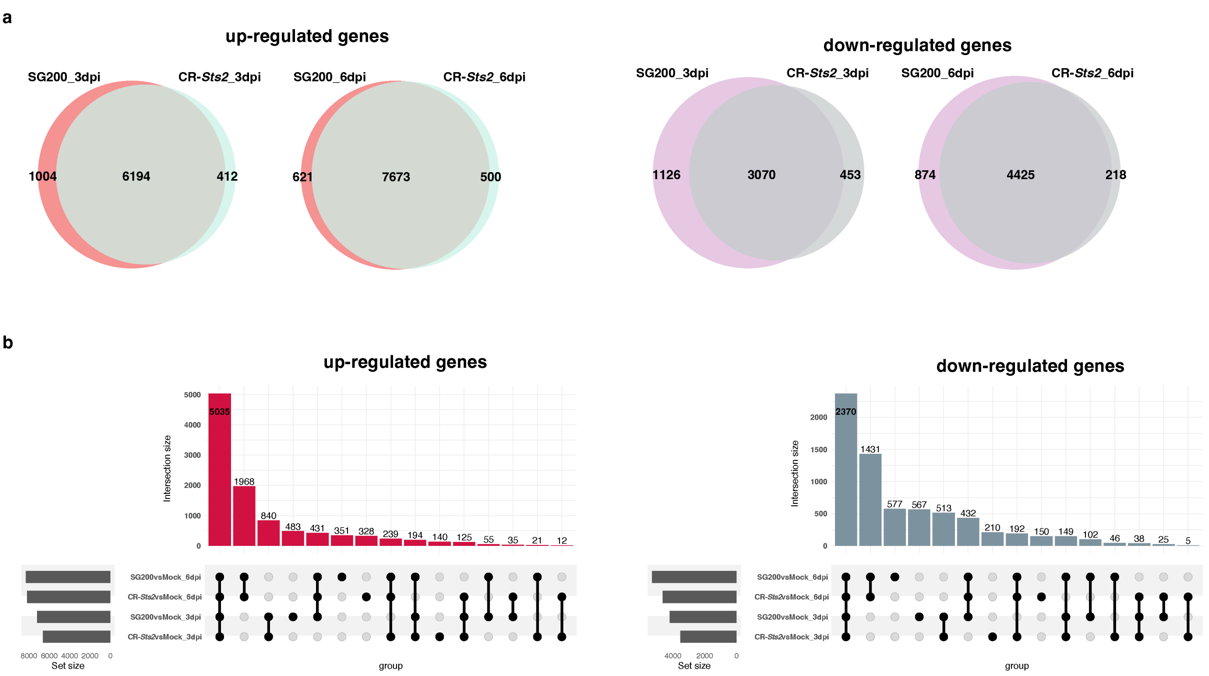


**Supplementary Fig3. Differentially expressed maize genes between *U. maydis* SG200 vs mock. a,** The Venn diagrams show the numbers of differentially expression genes between SG200vs. Mock and CR-Sts2 vs. Mock at 3 and 6 dpi, respectively. **b,** The intersection diagrams show the gene numbers in different paired comparison.

### Figure S4

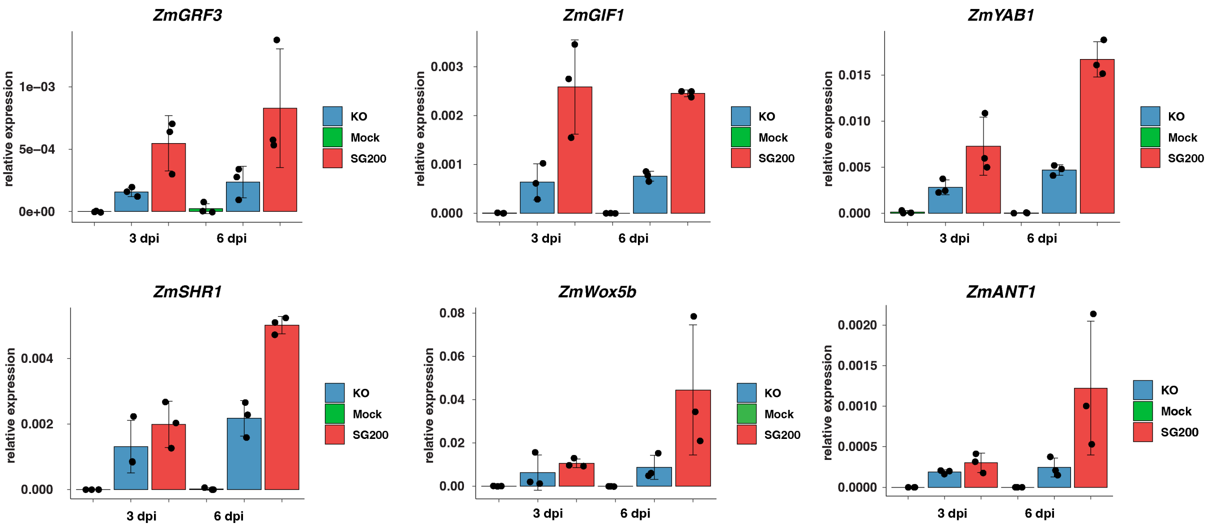


**Supplementary Fig4. The expression of maize genes upon *U. maydis* infection in RNA-seq samples by qPCR.** The expression level of leaf developmental regulators detected by qPCR in RNA-seq samples. The value of $2^{-\Delta Ct}$ between gene of interest (GOI) and *ZmGAPDH* are calculated and plotted.

### Figure S5

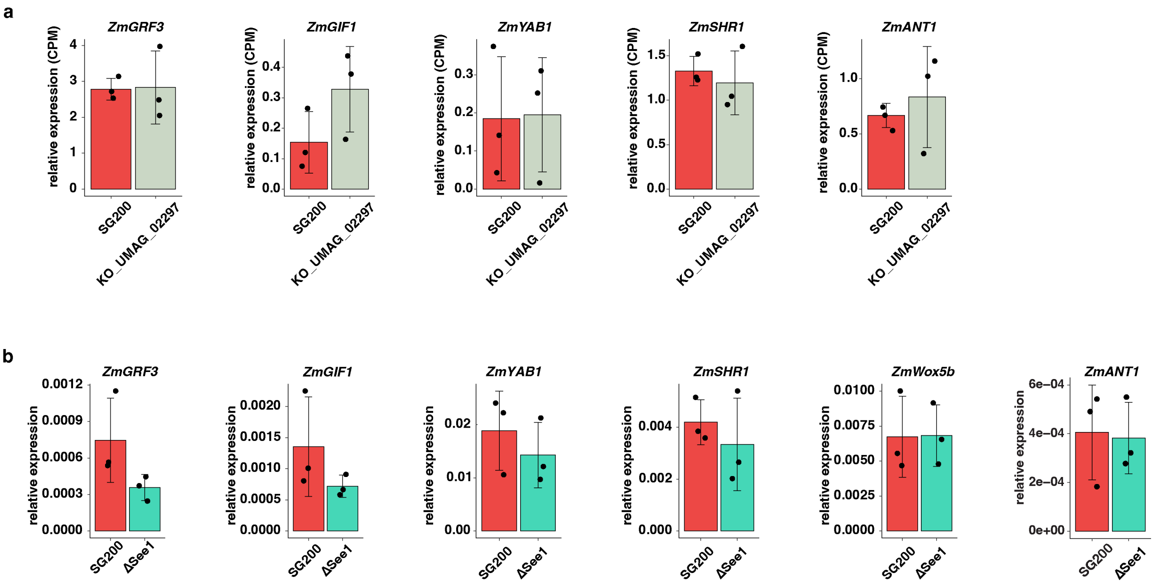


**Supplementary Fig5. The expression of maize genes upon different *U. maydis* effector mutants' infection. a,** The relative expression levels of maize leaf developmental regulators upon SG200 and KO_UMAG_02297 infection in maize line CML322 at 3 dpi from published data[37]. The CPM (counts per million) are plotted. **b,** The expression of leaf developmental regulators in SG200 and ΔSee1 mutant infected Golden Bantam at 6dpi by qPCR. The values of $2^{-\Delta Ct}$ between GOI and *ZmGAPDH* are calculated and plotted.
